# Theta-gamma coupling: a nonlinear dynamical model

**DOI:** 10.1101/304238

**Authors:** Alex Sheremet, Yuchen Zhou, Jack P. Kennedy, Yu Qin, Sara N. Burke, Andrew P. Maurer

**Affiliations:** Engineering School of Sustainable Infrastructure & Environment (ESSIE), University of Florida, 365 Weil Hall, Gainesville, FL 32611; Department of Neuroscience, McKnight Brain Institute, College of Medicine, University of Florida, P.O. Box 100244, 1149 Newell Drive, RML 1-100G, Gainesville, FL 32610; Department of Biomedical Engineering, University of Florida, Gainesville, FL 32611

**Author notes:** For correspondence (FMS); (FS). These authors contributed equally to this work (Alex Sheremet, Andrew P. Maurer).

## Abstract

Cross-frequency coupling in the hippocampus has been hypothesized to support higher-cognition functions. While gamma modulation by theta is widely accepted, evidence of phase-coupling between the two frequency components is so far unconvincing. Our observations show that theta and gamma energy increases with rat speed, while the overall nonlinearity of the LFP trace also increases, suggesting that energy flow is fundamental for hippocampal dynamics. This contradicts current representations based on the Kuramoto phase model. Therefore, we propose a new approach, based on the three-wave equation, a universally-valid nonlinear-physics paradigm that synthesizes the effects of leading order, quadratic nonlinearity. The paradigm identifies bispectral analysis as the natural tool for investigating LFP cross-frequency coupling. Our results confirm the effectiveness of the approach by showing unambiguous coupling between theta and gamma. Bispectra features agree with predictions of the three-wave model, supporting the conclusion that cross-frequency coupling is a manifestation of nonlinear energy transfers.

## Introduction

Hebb’s hypothesis that no psychological function can exist within any segment of cortex by itself (Hebb, 1958) has the profound implication that cognition emerges from the coordination of activity across the entire brain. However, if the brain works as a highly integrated device, with all parts working in concert, determining the mechanism of integration has proved a challenge so far (Lashley, 1958; Allen and Collins, 2013)

Because the most accessible observational data comes in the form of time series electrophysi-ological recordings, much of the effort of detecting integration has focused on the temporal scales of local-field potential (LFP). The temporal scales in extracellular recordings from the hippocampus range from 0.05 Hz to 500 Hz (Buzsaki and Draguhn, 2004), with readily identifiable features. The high end of the frequency spectrum is occupied by action potentials, which represent the activity of individual neurons and provide the “atomic” constituents of activity (McNaughton et al., 1996; Buzsaki and Draguhn, 2004; Buzsaki, 2006; Hasselmo, 2015; Eichenbaum, 2017; Buzsaki and Draguhn, 2004; Buzsaki, 2006). During awake behavior, the low-frequency end of the spectrum is dominated by theta oscillations (4-12 Hz), theoretically entraining global networks (Green and Arduini, 1954; Green and Machne, 1955; Vanderwolf, 1969; Lisman and Idiart, 1995; Buzsaki, 2002), and perhaps provide the temporal structure for the organization of intermediate scale oscillations such as gamma (broadly defined as 30-150 Hz; Buzsaki et al. 1983; Bragin et al. 1995; Belluscio et al. 2012; Lasztoczi and Klausberger 2014; Schomburg et al. 2014).

Efforts to detect how activity integrates across scales, however, have produced mixed results. On the one hand, the correlation of gamma amplitudes and theta phase is well established and interpreted as the organization of coactive neurons into cogent sequences in support of memory encoding and recall (Lisman and Idiart, 1995; DeCoteau et al., 2007; Tort et al., 2008,2009; Belluscio et al., 2012; Bieri et al., 2014). On the other hand, recent studies (Aru et al., 2015; **?**) warning of spurious results produced by observations of *n : m* phase coupling of theta and gamma cast doubts on the validity of previously reported phase-coupling observations (Belluscio et al., 2012; Zheng and Zhang, 2013; Xu et al., 2013,2015; Zheng et al., 2015).

Ambiguous phase-coupling results could be caused by the data analysis methodology stemming from current theoretical assumptions. The validity of any estimator is circumscribed by the validity of its underlying model. While there is no consensus on the mechanism of cross-frequency coupling, the Kuramoto model Kuramoto (1975) is commonly used as the paradigm for describing the phase evolution of biological oscillators, such as population emergence of cicadas, the synchrony of fireflies, entrainment of pacemaker cardiac cells, and neurons. The generic form of the model can be written as (e.g., Ermentrout 1998; Ermentrout and Kleinfeld 2001)

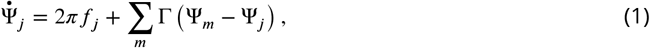

where Ψ_*j*_(*t*) is the phase of oscillator *j*, and the function Γ(Ψ_*m*_ – Ψ_*j*_) describes the interaction between the pair (*j,m*) of oscillators, and 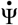 is the time derivative of Ψ. The phase Ψ is periodic in time, i.e., Ψ_*j*_(*t* + T_*j*_) = Ψ(*t*), where T_*j*_ is the period of the oscillator and 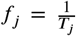. The sum only includes pairs of oscillators that are directly connected, therefore Γ implicitly defines the architecture of the network (Ermentrout and Kleinfeld, 2001). Equation 1 captures the basic dynamics of phase models (Cross and Hohenberg, 1993; Cross and Greenside, 2009). It describes the weak interaction of independent nonlinear oscillators, under two fundamental assumptions: 1) that the each oscillator evolves on an asymptotically stable limit cycle, and 2) that the leading order effect of weak interactions is a change in phase(Winfree, 1967; Kuramoto, 1984; Winfree, 2001). If an independent oscillator is represented in the phase space as a point moving on a closed path, the path itself can be described as relatively “stiff” under weak interactions^1^. Interactions only change the “speed” of the motion along the path, i.e., to introduce a phase shift. As phase shifts do not affect the total energy of the oscillator, such interactions necessarily involve only negligible energy exchanges between the oscillators.

While the model of a “stiff” limit cycle is theoretically suited for describing individual neurons as weakly-interacting oscillators, it might be less adequate for describing the dynamics of the Fourier components of hippocampal LFP. Recent studies show that, rather than being generated by well-defined, independent oscillators (such as individual neurons or groups of neurons forming “hardwired” neural circuits), hippocampal LFP frequencies represent propagating waves, i.e., transient neuronal populations co-active in a manner more akin to a metachronal rhythm (Lubenov and Siapas, 2009; Patel et al., 2012,2013; Petsche and Stumpf, 1960). Moreover, the strong response of gamma rhythms (factors of up to 5 in energy) to variations of theta energy (factors up to 100), as seen for example in the dependency of hippocampal activity to rat running speed (Whishaw and Vanderwol, 1973; Morris and Hagan, 1983; Chen et al., 2011; Ahmed and Mehta, 2012; Kemere et al., 2013; Zheng et al., 2015) strongly suggests that significant energy exchanges, and the energy flow through the brain should play a major part in the cross-scale coupling/integration process.

Here, we propose to use the alternative route of amplitude modeling (e.g., Cross and Hohenberg 1993; Passot and Newell 1994; Cross and Greenside 2009) which allows for simultaneous description of both amplitude and phase evolution. We hypothesize that hippocampal LFP rhythms represent waves of spontaneous collective neural activity that propagate across a two dimensional neural network of densely and randomly connected neurons. As the hippocampal LFP is primarily generated by synaptic transmembrane currents (Buzsaki et al., 2012), interactions among populations of neurons can be represented by Fourier modes that interact weakly, with significant effects on both the phases and the amplitudes of the interacting modes.This allows for the introduction of a simplified model that represents the basic elements of nonlinear interaction, the three-wave interaction model. The three-wave model is universal, in the sense that it characterizes the evolution of a large class of physical systems driven in the leading order by quadratic nonlinearity, in the same way the Kuramoto model (Kuramoto, 1975) can be said to capture the essence of phase models. A large body of literature is available that investigates the relevance and dynamics of three-wave interactions in a wide range of physical phenomena, including plasma (Coppi et al., 1969; Weiland and Wilhelmsson, 1977; Craick, 1985), nonlinear optics (Ablowitz and Segur, 1981; Boyd, 2003), internal oceanic waves (Phillips, 1977; Craick, 1985), and others.

Adopting the three-wave interaction model leads to a qualitatively different estimator for cross-scale coupling, the bispectrum, which is the leading order estimator in the well studied and understood sequence of higher-order correlators (cumulants) and their spectral representation (Hasselmann et al., 1963; Kim and Powers, 1965; Rosenblatt and Van Ness, 1965; Schreier and Scharf, 2006). The modeling is complimented by analyses of hippocampal LFP from freely-behaving animals. We present linear spectral analysis results that suggest the need of a general amplitude-phase nonlinear model. The three-wave model is introduced briefly, followed by a presentation of nonlinear analysis and a discussion of its general relevance for understanding the integration of brain activity in the service of cognition. Details data collection and the generic derivation of the three-wave interaction model conclude this study.

## Results

### Subjects and Behavioral Training

A total of four 4-9 months old Fisher344-Brown Norway Rats were used in the present study (Taconic), This was a mixed sex cohort comprised of 3 males (rats: 530,538 and 539) and 2 female (rats: 544, and 695) to incorporate sex a biological variable and begin to alleviate the disparity in research focused exclusively on males (Clayton, 2016). In the present study, we had no a priori predictions that sex will alter oscillations of the hippocampus as single-unit physiology is relatively consistent in female animals across estrous (Tropp et al., 2005). Upon arrival, rats were allowed to acclimate to the colony room for one week. The rats were housed individually and maintained on a reverse 12:12 light/dark cycle. All training sessions and electrophysiological recordings took place during the dark phase of the rats’ light/dark cycle. Training consisted of shaping the rats to traverse a circular track or figure-8 maze for food reward (45mg, unflavored dustless precision pellets; BioServ, New Jersey; Product #F0021). During this time, their body weight was slowly reduced to 85% to their ad libitum baseline. Once the rat reliably performed more than one lap per minute, they were implanted with a custom single shank silicon probe from NeuroNexus (Ann Arbor, MI). This probe was designed such that thirty-two recording sites, each with a recording area of 12 by 15 μm, were spaced 60 μm apart allowing incremental recording across the hippocampal lamina. In preparation for surgery, the probe was cleaned in a 4% dilution of Contrad detergent (Decon Contrad 70 Liquid Detergent, Fisher Scientific) at 55°C for 2 hours and then rinsed in distilled water (Vandecasteele et al., 2012).

All behavioral procedures were performed in accordance with the National Institutes of Health guidelines for rodents and with protocols approved by the University of Florida Institutional Animal Care and Use Committee.

### Neurophysiology

Following recovery from surgery, the rat was retrained to run unidirectionally on a circle track (outer diameter: 115 cm, inner diameter: 88 cm), receiving food reward at a single location. Following a few days of circle track running, the rats were trained to run on a digital-8 maze (121×101cm, LxW). During these sessions, the local-field potential was record on a Tucker-Davis Neurophysiology System (Alachua, FL) at ∼24k Hz (PZ2 and RZ2, Tucker-Davis Technologies). The animals position was recorded at 30 frames/s with a spatial resolution less than 0.5 cm/pixel.

### Spectral evolution as a function of rat speed

As discussed in the introduction, phase-coupling estimators derived from phase-model paradigms (e.g., **?**) implicitly ignore variations of amplitude (energy). However, recent studies reported increases in gamma frequency and power with rat speed (Chen et al.,2011; Ahmed and Mehta, 2012; Kemere et al., 2013; Zheng et al., 2015). During awake-exploration, the hippocampal LFP exhibits coordinated patterns of theta and gamma (Bragin et al., 1995; Chorbak and Buzsaki, 1998). In rats, the amplitude and frequency of the theta rhythm increase with running speed (Whishaw and Vanderwol, 1973; Morris and Hagan, 1983), which also correlates with an increase in gamma frequency and power (Chen et al., 2011; Ahmed and Mehta, 2012; Kemere et al., 2013; Zheng et al., 2015). As firing rates of both the hippocampal and entorhinal neurons increase with running speed (McNaughton et al., 1983; Rivas et al., 1996; Shen et al., 1997; Czurko et al., 1999; Hirase et al., 1999; Maurer et al., 2005; Kropff et al., 2015), it can be inferred that energy into the hippocampus increases with animal speed. This suggests that the evolution of energy distribution over scales, and implicitly the flow of energy through the neural system, play an important part of the brain integration process, and the development of cross-scale coupling.

Although previous work examined variance statistics (Fourier or wavelet spectral analysis), to our knowledge, a systematic examination of the dependency of LFP power spectral density on speed and hippocampal layer has not been conducted. Figure 1 shows the spectral density in the str. pyramidale (CA1.pyr), radiatum (CA1.rad), lacunosum moleculare (LM) and upper blade of the dentate gyrus (DG) as a function of speed. Our observations of the power spectra can be characterized in general as having a power law *f^−α^* structure. However, unlike previous reports (Bak et al., 1988; Beggs and Plenz, 2003; Buzsaki, 2006; Ahmed and Mehta, 2012; Gao, 2016), the spectral slope a is not constant throughout the entire frequency range. Instead, at least two domains with distinct slopes can be identified, most evident in the LM and DG layers. The domains are consistent across layers and form rather well-defined frequency intervals that can be associated with theta and its harmonics (Sheremet et al., 2016) and gamma (∼60-130 Hz). At high speed, the DG theta shows clearly distinguishable harmonics at 16Hz, 24Hz, 32Hz, and a final, noticeable peak at 40Hz (data from `r539♀-maurer`; further validation of multiple theta harmonics provided by nonlinear analysis below). Adjacent hippocampal layers also exhibit harmonics of theta, although not as prominent as the DG region. In agreement with previous results (Bragin et al., 1995), theta and gamma power are higher in the LM and DG than the CA1 layers. Observed gamma activity supports the hypothesis that, as the hippocampus receives more speed related input, activity is captured among the recurrent local circuits that support gamma resulting in higher power in these frequencies.

**Figure 1.**
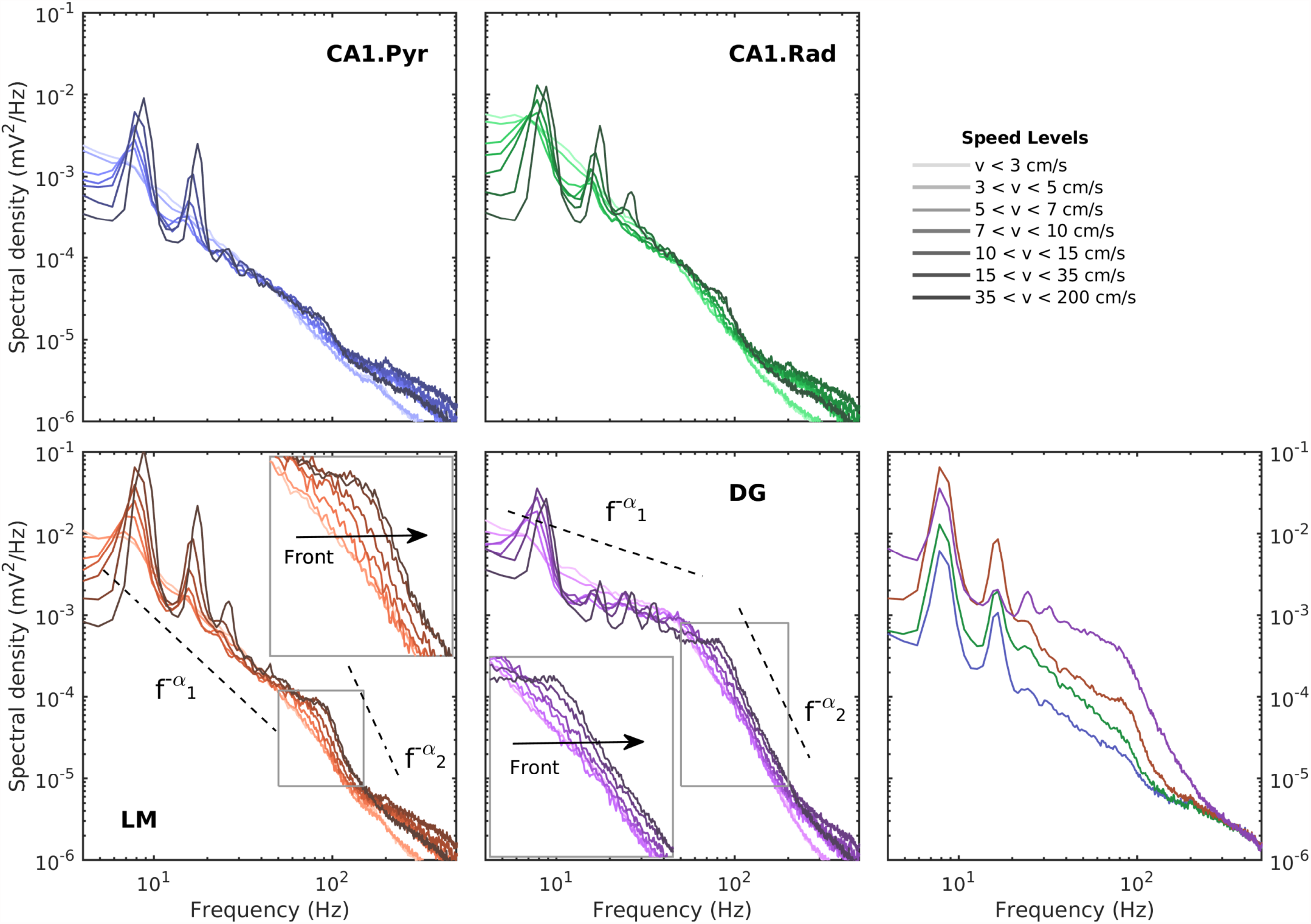
Changes in power spectral density as a function of velocity and layer. Left and middle: Comparison of the evolution of the power spectrum in the CA1.pyr and CA1.rad layers as a function of speed (transition from lighter to darker hues indicate low– to high velocities). The increases in theta and theta harmonic with speed are associated with a mild increase in gamma power. Insets show details of the evolution of the gamma range (magnification of gray rectangle area). Spectral shapes in all layers examined follow a power laws of the type *f^−α^*. Remarkably, in the LM and DG layers, the frequency domains of theta (and its harmonics) and gamma exhibit obviously distinct slopes. In the LM and DG regions, as power increases in the gamma range, the spectrum preserves its slope and shifts to the right, creating the appearance of a movoing front. Right: Comparison of power spectral densities across layers at the highest speed bin. Note that all regions exhibit multiple harmonics of theta. In agreement with prior publications (Bragin et al., 1995), the CA1.pyr layer shows the least energetic gamma range of all the layers examined. Data produced from rat `r539♀-maurer`.

In the 50-130 Hz frequency range, the evolution with speed of the distribution of energy acquires the form of a front of constant slope shifting to the right, toward higher frequencies (insets in figure 1). This effect is observed across all layers, although most prominent in the DG. This strongly suggests that, as rat speed increases, a weak nonlinear coupling of hippocampal activity develops across the spectrum, leading to a weak mean flow of energy from low to high frequencies. In line with this idea, the organization of spike times across assemblies of neurons in the hippocampus has been equated to the organization of weakly connected, quasi-periodic oscillators (Izhikevich, 1999b,a). Specifically, ion channels expression endows hippocampal pyramidal neurons with intrinsic, voltage-dependent oscillations near theta frequency theoretically leading to potential entrainment and organization when driven by an external input (Leung and Yim, 1991; Kamondi et al., 1998). Furthermore, it has been suggested that CA1 and CA3 networks behave akin to voltage-controlled oscillators with unique voltage/resonance properties (Sullivan et al., 2011). Theoretically, as the excitation increases within the CA1 and CA3 regions, the resonance of the system transitions from a basal frequency towards higher frequencies (Sullivan et al., 2011). From this insight, it can be inferred that, as theta power increases as a function of speed, there is a direct cascade of the energy towards modulating the resonance of networks in the gamma frequency. The hypothesis of nonlinear energy transfers (i.e., nonlinear cross-frequency coupling) developing as rat activity increases suggests the remarkable consequence that theta-gamma phase-coupling (if any) develops during this evolution and should also depend on speed.

This brings back the question of the methodology of detecting cross-frequency phase correlations in LFP Traces. Following the skepticism expressed in the study by **?**, and in light of the above results, we propose to set aside the phase model approach and seek an alternative nonlinear formulation that incorporates both amplitude and phase coupling (amplitude models, e.g., Cross and Hohenberg 1993; Cross and Greenside 2009). Such models are ubiquitous in physics (Zakharov et al., 1992; Nazarenko, 2011; L’vov, 1998).

### The three-wave equation: a nonlinear amplitude model paradigm

#### The dynamical equation

Here, we summarize the ideas behind the derivation of the amplitude equation and the simplified three-wave equation. Some details of the algebra are given in the Appendix. Following the studies of Lubenov and Siapas (2009), Patel et al. (2012), and others, we postulate that the neural activity in the hippocampus can be represented as a multiple-scale, weakly-nonlinear wave field, whose dynamics is governed by nonlinear interaction. For simplicity, we limit the discussion to a one-dimensional spatial system whose small deviation from an equilibrium state is defined by a function *εφ*(*x,t*), e.g., electric potential, of spatial coordinate vector *x* and time *t*. Under quite general conditions, the Fourier components of *φ*

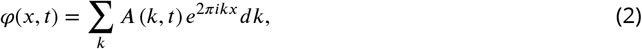

satisfy a dynamical equation of the form

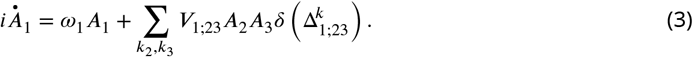

Here *k* = 1/*λ* is the wave number (wavelength *λ*), and *A_j_* = *A* (*k_j_,t*), *ω_j_* = *ω*(*k_j_*) = 2*πf*(*k_j_*), are the complex amplitude and radian frequency of mode *j*, the latter given by the dispersion relation 23, and we used the shorthand notation 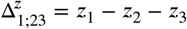(e.g., *k* in equation 27).

Equation 3 can be interpreted as a general model for the nonlinear evolution of the different LFP rhythms, identified by their wave numbers *k*, and representing different “scales” of the system. The right-hand side sum represents the nonlinear interaction of mode *k*_1_ with pairs of wave numbers (*k*_2_,*k*_3_). A triplet of interacting modes (*k*_1_,*k*_2_,*k*_3_) is usually referred to as a triad. The interaction coefficient *V*_1;23_ is a function of the wave numbers k*j*, *j* = 1,2,3. An important element, introduced by the orthogonality of the Fourier modes (equation 21, Appendix) is the factor 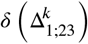, which acts as a selection criterion: only the triads that satisfy the equation

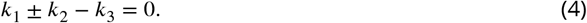

are counted in the sum in equation 3.

Note that the wave-number representation is different than the frequency one used, e.g., in figure 1; however, the dispersion relation *f* = *f(k)* (equation 23, Appendix) readily allows for switching between the two. In the rest of this paper, we will treat the concepts of “wave number”, “mode” and “scale” as equivalent and interchangeable. Hence, the dynamical equation 3 describes the full effects of cross scale interaction, including cross-scale energy transfers (expressed in amplitude variations), and synchronization effects (expressed in phase coupling), under the assumption of weak interaction.

#### The three-wave equation

Although the dynamical equation 3 should be solved for the full spectrum of *k*, much can be learned from studying the interaction of a small number of interacting modes. For quadratic nonlinearities the smallest such subset is a single triad, a triplet of modes *k_j_*, *j* = 0,1,2, satisfying the selection criterion *k*_1_ ± *k*_2_ − *k*_3_ = 0. Restricting equation 3 to a single triad, after a scaling change of functions (see Appendix) the equation acquires the canonical form

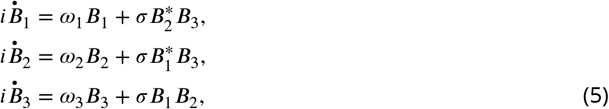

where *σ* is the interaction coefficient, and *B_j_* are new, scaled, modal amplitudes. Equation 5 is typically referred to as the three-wave equation (Craick, 1985; Rabinovich and Trubetskov, 1989). A large body of literature is available that investigates the relevance and dynamics of single-triad interactions in many physical situations, including plasma physics (Coppi et al., 1969; Weiland and Wilhelmsson, 1977; Craick, 1985), nonlinear optics (Ablowitz and Segur, 1981; Boyd, 2003), internal oceanic waves (Phillips, 1977; Craick, 1985), and other fields. The meaning of equation 5 becomes evident when written in alternative forms.

For the cross-frequency coupling discussed in this study, single triad interactions are relevant for: a) theta-gamma-gamma interactions, where *k*_1_ represents a theta mode, and *k*_2,3_. are gamma modes, with *k*_1_ ≪ *k*_2,3_, and b) theta-theta-theta harmonic, where *k*_1_ ≈ *k*_2_, and *k*_3_=*k*_1_ + *k*_2_, where *k*_l,2_, are both theta modes and *k*_3_ is the harmonic of theta (figure 2).

**Figure 2.**
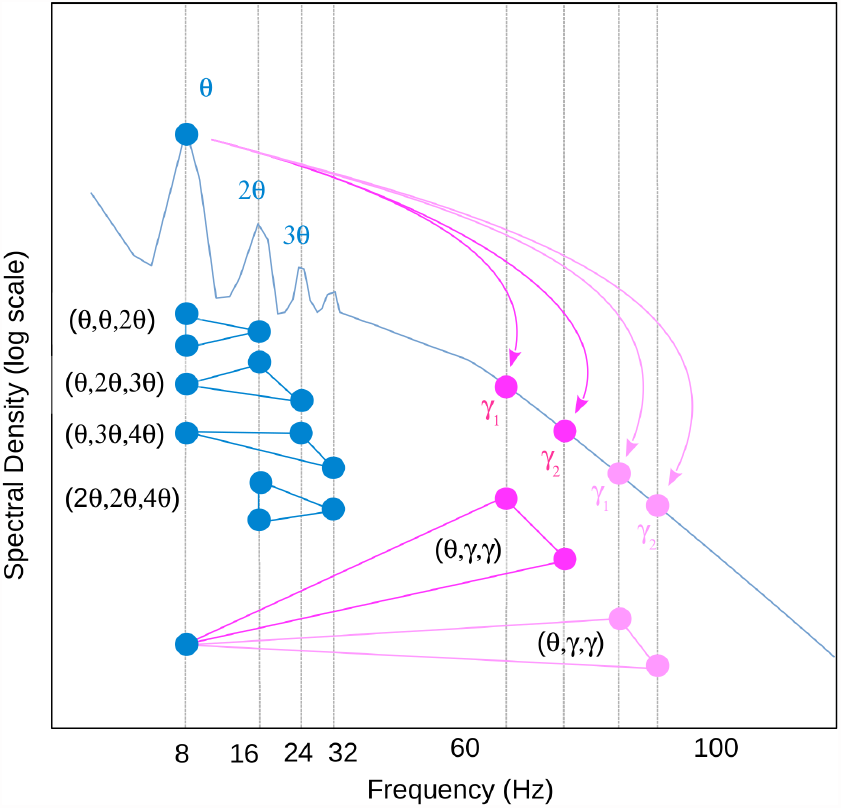
Schematic of different types of interacting triads (equations 3) belonging the theta-harmonics family (blue) and theta-gamma family (magenta).

#### Phase-amplitude and energy evolution

Switching to amplitude-phase representation *B_j_* = *b_j_e^iΘ_j_^*, with *b_j_*, 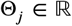 and *b_j_* > 0, equations 28 become

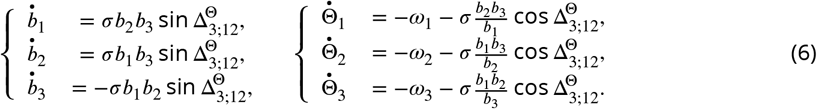

This form highlights the effect of the interaction on modal amplitude and phases. The following statements should be obvious. Cross-scale nonlinear coupling modifies both energy (amplitude) at a specific scale, and the phases of the interacting scales. Nonlinear effects depend on *the magnitude of the modal amplitudes (modal energy)*: increasing the energy into the system amplifies nonlinear effects. The effect of the nonlinearities is a redistribution of energy across interacting modes. Synchronization effects (phase coupling) is stronger for a mode that has lower energy. The coupling between the three interacting modes is measured by the statistics of the triple phase 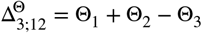

Alternatively, some straightforward algebra yields for modal amplitude squared the equations

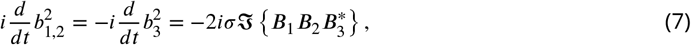

where 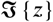 is the imaginary part of the complex number *z*. Note that the triple product 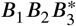 incorporates information about both modal amplitudes (squared), and phases (through the phase difference 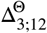). Averaging equations 7 yields

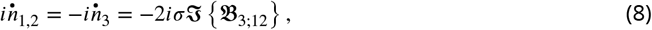

where E is the expected value operator, and

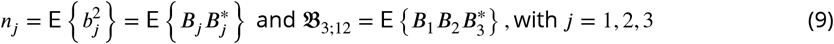

are recognized to be the first two leading order amplitude correlators. The second order correlator *n_j_* is the variance of mode *j* (loosely referred to as modal energy, although the units might not be energy units), and is proportional to the spectral density (e.g., figure 1). The third order correlator 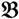 is called *bispectrum*.

It should be clear now that the three-wave model (equation 5) and the Kuramoto phase model (equation 1) describe different physical phenomena: the phase model describes the interaction of oscillators that represent physical entities, such as neurons or well-defined neuronal circuits; the three-wave model represents interacting transient assemblies of neurons. In the phase model, there is no exchange of energy between oscillators (no “amplitude” variation); in contrast, amplitudes and their evolution play a central role in the three-wave model. In the phase model, the smallest interacting group of elements is composed of *two* oscillators; in the three-wave model, the smallest group is a *triad – three* modes. The selection of interacting elements is also different: in the phase model the oscillators have to be physically connected, while in the three-wave model the mode selection is based on mutual forcing of overlapping waves with matching wave numbers (see equation 4).

The fundamental differences between the two models have significant consequences on the methodology of data analysis. Equations 6 and 8 highlight the importance of the bispectrum 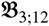 as a measure the strength of the nonlinear interaction and synchronization. The real and imaginary part of the bispectrum is related to third-order statistics of the entire LFP (including all frequency bands), through

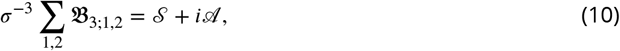

where 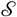 is the skewness and 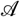 is the asymmetry, and *σ* is its standard deviation (Masuda and Kuo, 1981; Elgar, 1987). The normalized bispectrum (Haubrich and MacKenzie, 1965; Masuda and Kuo, 1981; Elgar, 1987; Sheremet et al., 2016, 2017)

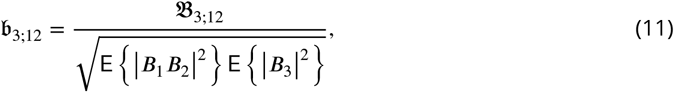

eliminates the distortion induced by the frequency distribution of variance, and has the convenient property that 0 ≤ 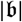 ≤ 1. The modulus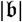 is called *bicoherence* and the angle arg 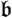 is called *biphase*.

Because the bicoherence is statistically zero if modes *j* = 1,2,3 are statistically independent, it is commonly used as an “index” of nonlinearity strength. The biphase provides a measure of the average phase difference 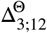 (see equations 6). In the sequel, we discuss the insights provided by the bispectral analysis of the LFP frequency content.

### Cross-frequency coupling: evidence of theta-gamma interaction

The three-wave interaction paradigm highlights the important role of the evolution of energy in the generation of cross-frequency coupling. It also points to the normalized bispectrum 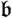 (equation 11) as an adequate statistical estimator for the strength of coupling and average phase coupling. Because the rat speed is a strong indicator of the energy into the system (figure 1), bispectral estimates are also ordered as a function of rat speed. The interpretation of bispectral plots is discussed in Sheremet et al. (2016) and many other places. Here, we just note that the position (*f*_1_, *f*_3_) of statistically significant bicoherence value (zero-mean bicoherence 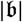 < 0.1 at 95% confidence level) indicates an interacting triad of modes (*f*_1_, *f*_2_, *f*_1_ + *f*_2_).

As previously reported (Sheremet et al., 2016), the bispectral estimates also show significant variability with rat speed. Low-speed bispectra (*v* < 10 cm/s Fig. 3; data from `r539♂-maurer`) show barely-significant phase coupling. The activity in the CA1 layer is nearly-linear. The LM and DG layers show weak coupling between theta and its first harmonic. A weak coupling between theta and gamma is also seen in the DG.

**Figure 3.**
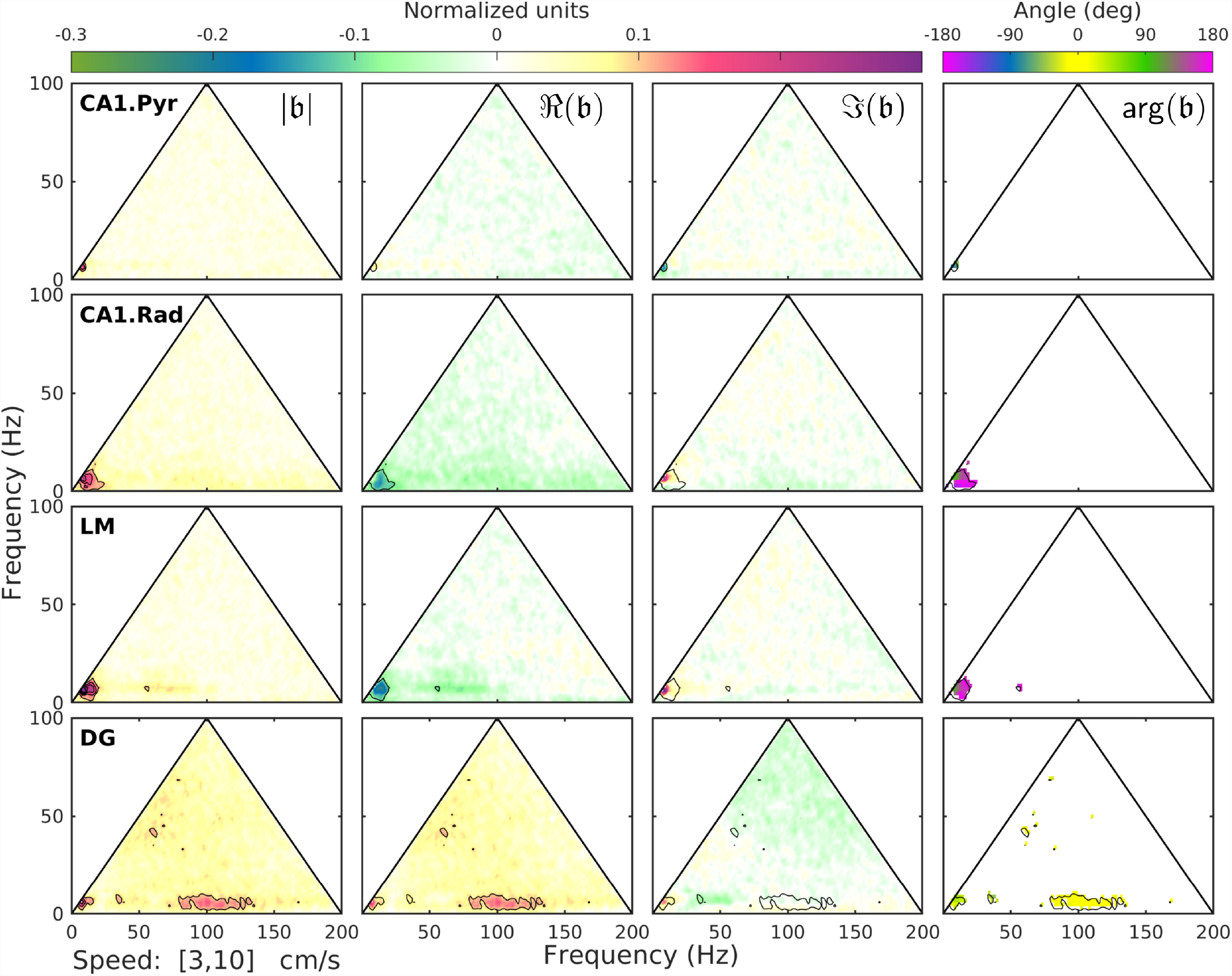
Normalized bispectrum (Equation 11) at low speeds (*v* < 10 cm/s). An in depth explanation of the bicoherence plot can be found in Sheremet et al. (2016). Briefly, the triangular region represents the area containing non-redundant information for the discrete Fourier transform. Columns (left to right) show the bicoherence 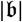, the real 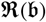 and imaginary 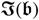 parts of the normalized bispectrum 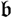, and the biphase arg(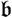). The first three can be interpreted as nonlinearity strength, and measures of the contribution of different triads to the skewness and asymmetry of the LFP. Peak in the bispectral estimate represents a phase-coupled triplet (*f*_1_, *f*_2_, *f*_1_ + *f*_2_), where, *f_j_*, *j* = 1,2 are frequency bands in the Fourier representation. The observations in each row correspond to a given hippocampal layer. For each layer, black contours mark the significant bicoherence value of 0.1 (with 300 DOF, zero-mean bicoherence is <0.1 at 95% confidence level). Data produced from rat r539_♂_-maurer.

In contrast, at higher running speeds (i.e., *v* > 40 cm/s, figure 4), the bispectral map show a rich phase-coupling structure involving theta and gamma triads. In agreement with the observations of spectral evolution (figure 1) and previously reported results (Sheremet et al., 2016), strong phase coupling develops between theta (theta frequency *f_θ_* ≈ 8 Hz) and its harmonics (*nf_θ_*, with *n* = 2,3,4,…), with bicoherence peaks detectable at frequencies below ∼50 Hz, at various strengths, across all layers. The strongest peaks are seen in the LM and DG regions, with interacting harmonics reaching as high as (5,1,6), (3,2,5), and (3,3,6). Note also that up to *n* = 5 harmonics can be identified in the DG spectrum (figure 1). The development of the theta harmonics strongly coupled to theta indicates a nonlinear deformation of the theta rhythm toward positive skewness and asymmetry (figure 4), and should not be attributed to the generation of additional, statistically independent rhythms (Sheremet et al., 2016). Remarkably, theta harmonics play different roles in the statistical deformation of the theta rhythm. For example, in the DG layer, triads (1,1,2), (2,1,3) generate positive skewness, 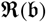 > 0, and weak positive asymmetry 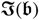; however, triads (3,1,4), (4,1,5) generate negative skewness, 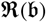 < 0 (figure 4).

**Figure 4.**
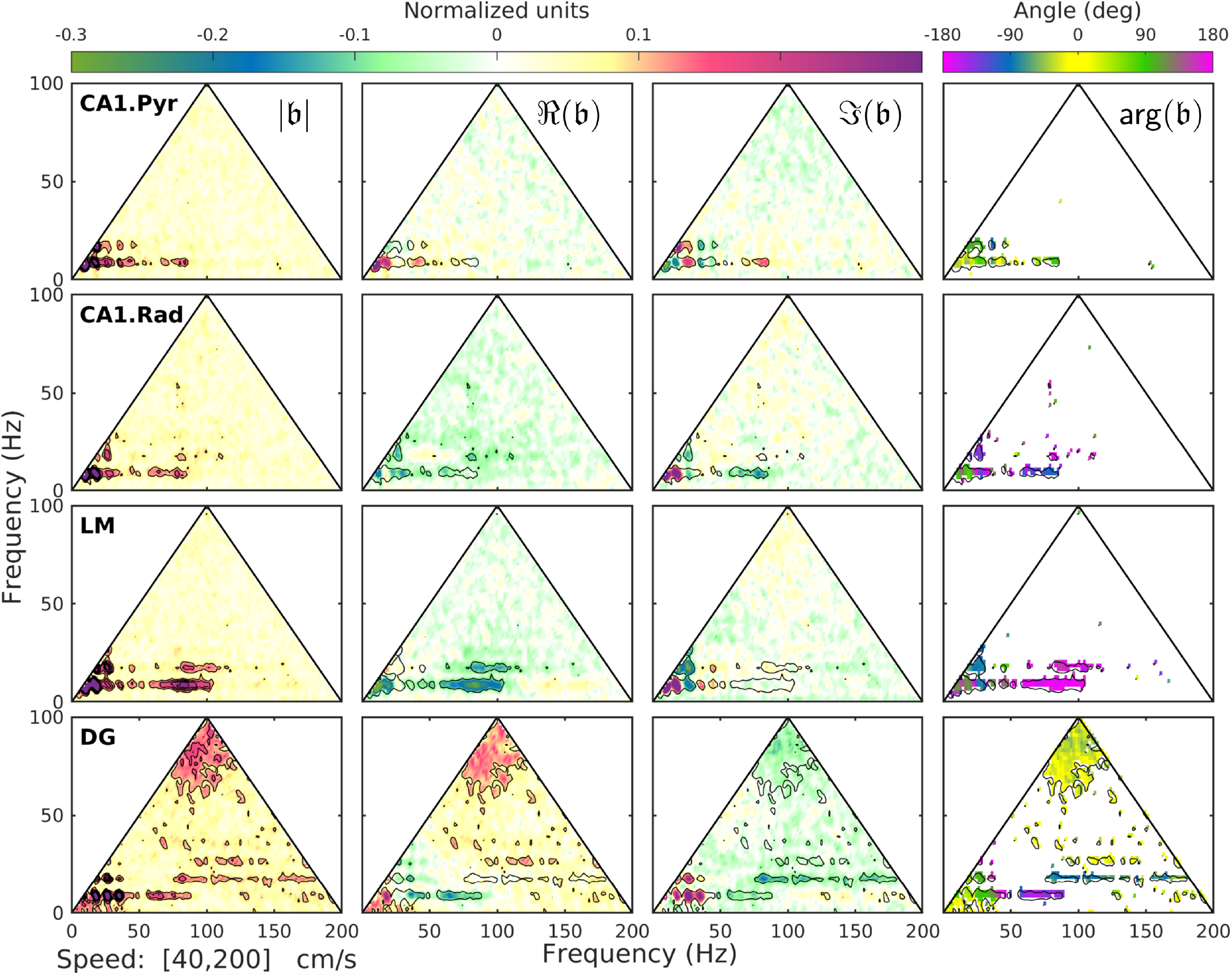
Normalized bispectrum at high speeds (*v* < 40 cm/s). Same organization as in figure 3. Note the development of multiple regions of high bicoherence 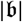, notably: 1) theta (*f_θ_* ≈ 8 Hz) and its harmonics (*nf_θ_*, with *n* = 2,3,4,…), reaching as high as (5, 1,6), (3,2,5), and (3,3,6) in the DG layer, and 2) theta and a wide gamma band spanning approximately the interval between 60 Hz to 100 Hz. The LM and DG layers show the richest nonlinear structures of the four layers examined. See the text for a detailed discussion of the results. Data produced from rat r539_♂_-maurer.

The increase of theta energy and nonlinearity is, however, accompanied by the development of significant phase coupling between theta and gamma, related to interaction between rhythms, rather than to nonlinear shape deformation. Theta-gamma coupling covers the entire gamma frequency band (the wide bands located approximately at 60 Hz < *f*_1_ < 100 Hz and *f*_2_ ≈ 8 Hz). The effect is again strongest in the LM and DG regions (see bicoherence plots in figure 4), where interacting triads of the form (*f_γ_*, *f_θ_*, *f_γ_* + *f_θ_*), and (*f_γ_*, 2*f_θ_*, *f_γ_* + 2*f_θ_*) are obvious, where *f_γ_* is a frequency in the gamma band. The overall effect of theta-gamma interaction on the time series is to generate negative skewness (see LM and DG layers, figure 4), with a biphase consistently ≈ 180°; the exception is the DG layer, where the biphase of triads of the type (*f_γ_*, 2*f_θ_*, *f_γ_* +2*f_θ_*) show biphase values of ≈ −90°. In turn, the gamma band develops a broad harmonic band in the 150-200 Hz band, corresponding to interacting triads of the type (*f_γ_*, *f_γ_*, 2*f_γ_*).

Because the location and magnitude of the bicoherence peaks identify the interacting triads and measure the intensity of their interactions, the strength of the nonlinear coupling can be quantified for each type of interaction by integrating the bicoherence over the region of interest (Sheremet et al. 2016; see e.g., figure 5). In agreement with the results presented in Sheremet et al. (2016), the nonlinearity is significant at faster running speeds (*F*[_1,16_] = 38.69, *p_T_* = 0.001; repeated-measures, where *p_T_* is Tukey’s range test, Tukey HSD). Moreover, the strength of nonlinearity also varies significantly as a function of layer (*F*[_3,16_] = 8.63, *p_T_* = 0.0001), with the LM being significantly more nonlinear than the CA1.pyr (*p_T_* = 0.23), and CA1.rad (*p_T_* = 0.002). The DG and LM regions show a similar degree of nonlinearity (*p_T_* = 0.80), significantly stronger than the CA1.rad (*p_T_* = 0.01). Speed appears to have a weaker effect on nonlinearity in the LM and DG relative to the CA1 layers (*F*[_3,16_] = 2.56, *p_T_* = 0.09)

**Figure 5.**
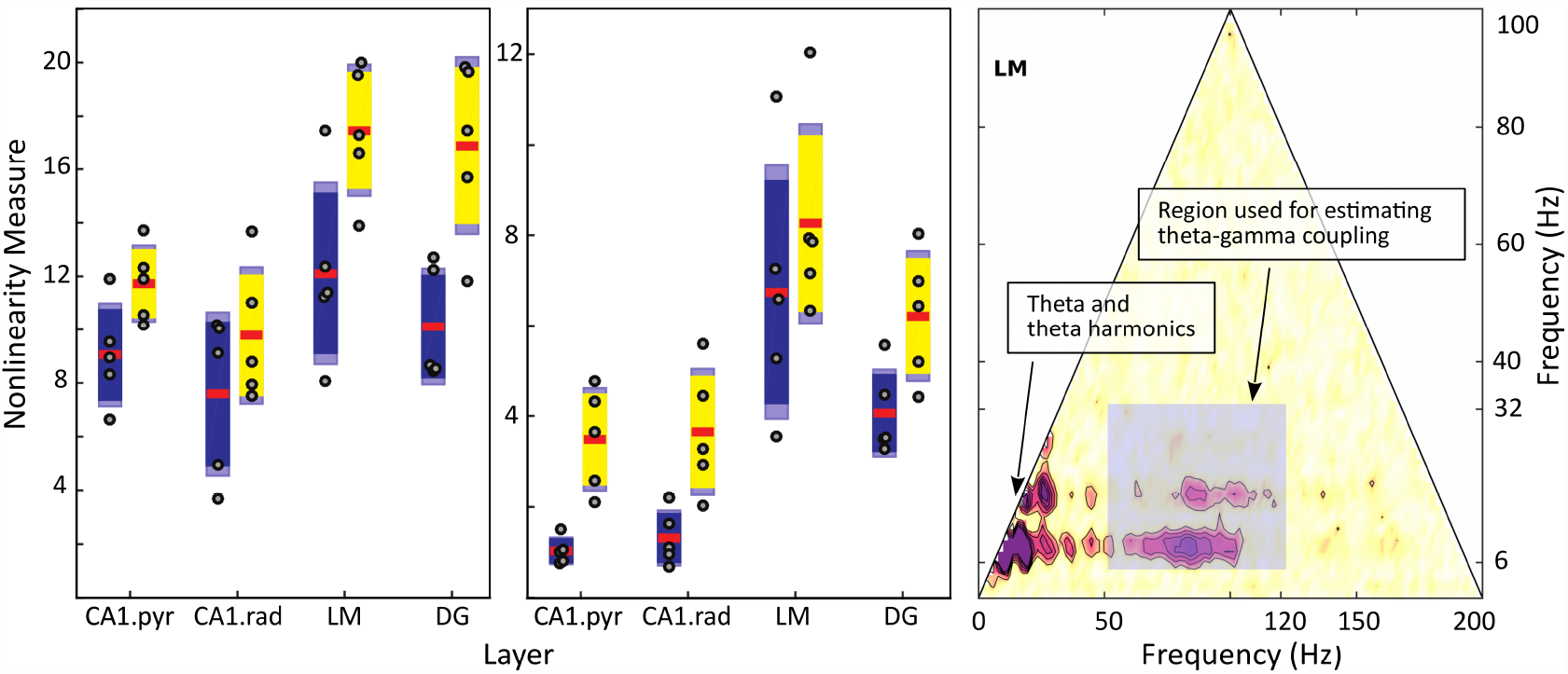
Estimate of nonlinear cross-frequency phase coupling. Left: Total overall strength of nonlinearity, estimated by summing the bicoherence over its entire definition domain. Estimates are shown for 5-15cm/s speed bin (blue) of >35cm/s speed bin (yellow). Middle: Strength of nonlinear coupling between theta and gamma rhythms estimated by summing the bicoherence values in the shaded rectangle (right panel). The LM and DG layers show significant increase in coupling strength compared with the CA1 layers (see also figures 3-4, and the discussion in the text). Both total and strictly theta-gamma coupling strength show significant variability as a function of rat speed and hippocampal layer. The theta range includes its harmonics. Note that the y-axis differs between plots in order to optimize for display of the theta-gamma-gamma nonlinearity. Red is the mean, solid dark blue or yellow is 1.96 of the standard error of the mean (S.E.M.) and light blue denotes the boundaries of 1 standard deviation. Right: Region used for estimating the strength of theta-gamma coupling is a rectangle covering the intervals [6 Hz, 32 Hz] for theta and [50 Hz, 120 Hz] for gamma.

The strength of nonlinearity in the area of interest for theta-gamma coupling (rectangle spanned by the theta and gamma bands [6 Hz, 32 Hz] × [50 Hz, 120 Hz]; figure 5) shows a strong effect of speed on nonlinearity (*F*[_1,16_] = 52.82, *p* = 0.0001). The strength of nonlinearity in the theta and gamma frequency ranges (*F*[_3,16_] = 15.40, *p* = 0.0001) depends significantly on the layer. Post hoc analysis indicated that the LM and DG were both significantly more nonlinear compared to the CA1 layers (*p* < 0.05). Finally, nonlinearity increase with running speed at similar rates across all layers (*F*[_3,16_] = 0.51, *p* = 0.59).

While the magnitude of the normalized bispectrum is easy to interpret and understand as an index of the nonlinearity strength, the meaning of the biphase is more opaque, even interpreted as an estimate of average phase difference 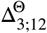, because it involves the sum of three phases. The estimates of the biphase in figure 4, however, show intriguing, well-defined structures. Although studying the biphase in the proper context requires a stochastic theory for the three-wave model, it should be possible to use the dynamical form of the three-wave model to derive some insight into the behavior of the system.

Of the rich structure of the biphase map, some prominent features are relatively easily explained. For example, in the DG layer (figure 4), the biphase for theta/harmonic coupling is ≈ 0, while triads involving theta and gamma have biphase values of ≈ 180°. It is obvious that a phase differences Δ^Θ^ values of 0 or 180° have a special meaning. Consider the three-wave equations written in the form given by equation 32, Appendix. Setting tentatively 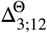 = 0 or 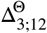 = *π* simplifies the system to

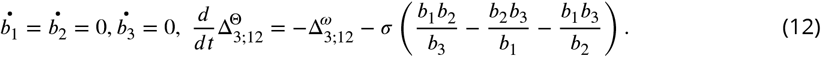

but the resulting biphase, the *average* of 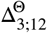, will not cancel in general because 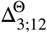 evolves due to frequency mismatch and the nonlinear interaction term. Recall that in equation 5 the modes are identified by their wave numbers, satisfying equation 4; however, whether 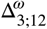 = 2*π* ([*f(k_3_* – *f(k_1_* – *f(k_2_*]) cancels or not depends on the form of the dispersion relation 23. To keep the phase difference constant the two terms in the last equation should cancel, i.e., if the frequencies should match in the same way as the wave numbers,

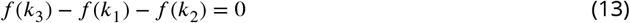

and the amplitudes squared should satisfy the equation

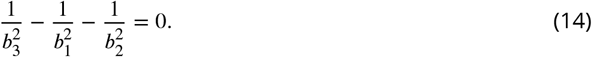

If these assumptions are met, the triad (*k_1_*, *k_2_k_3_*) is stationary.

Taken together, equations 4 and 13 are known as the resonance criterion (e.g., Craick 1985; Kartashova 2010), and play a key role in the long term dynamics and the stochastic theory of nonlinear systems. Equation 14 implies that the system achieves a stationary state only if the modal amplitudes are in a certain relation to each other. The obvious solution, 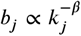, with 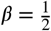 is significant for this discussion because it provides an intriguing explanation to the power-law shape of the spectra observed in the hippocampus (figure 1): if *k* = *f^α^* (see the dispersion relation 23, Appendix) the frequency spectrum has the tantalizing form *f^−α^*. This could also provide an explanation for the variations of the spectral slope seen in figure 1, if the dispersion relation has a scale dependency, as is common in wave physics (see e.g., **?**).

The effect of the harmonic of theta being on average in phase with theta is to sharpen the crests and flatten the troughs of the time series (positive skewness, see figure 6, left panel). This is corroborated by the real part of the bispectrum, 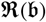 > 0, for the peak located at (*f_θ_*,*f_θ_*,*f_2θ_*) in the DG layer (figure 4). To understand the effect of the biphase value of 180° for theta-gamma coupling, consider three waves of the form (e.g., equation 25)

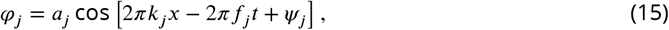

where *k*_1_ represents the theta mode, and *k*_2,3_, are gamma modes, with *k*_1_ = *k*_3_ – *k*_2_, *k*_1_ ≪ *k*_2,3_, and assume that the amplitudes of the gamma modes are constant, and the wave numbers and frequencies satisfy the resonance conditions 4 and 13. Recall the elementary discussion of the group structure generated by two oscillations with slightly different wave numbers and frequencies (as are the gamma modes): their superposition has the form

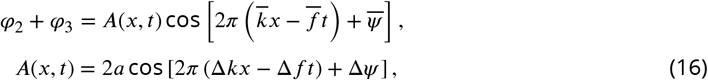

where *a* = *a*_1_ ≈ *a*_2_ is the amplitude of the gamma modes, 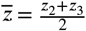, 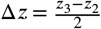, and where *z* is either *k*, *f*, or *ψ*. The gamma envelope *A* is modulated on the long temporal/spatial scales of Δ*k*^−1^ and Δ*f*^−1^. In this description, the biphase represents the average value of initial phase difference, i.e., on average, *ψ*_3_-*ψ*_2_-*ψ*_1_ = *π*. Without loss of generality, we can set *ψ*_1_ = 0 to obtain that, on average, *ψ*_3_ – *ψ*_2_ = 2Δ*ψ* = *π*; using the resonance conditions 4 and 13, we also have *f*_1_ = *f*_3_ – *f*_2_ = 2Δ*f*. Therefore, at *x* = 0, theta, and the gamma envelope are, respectively

**Figure 6.**
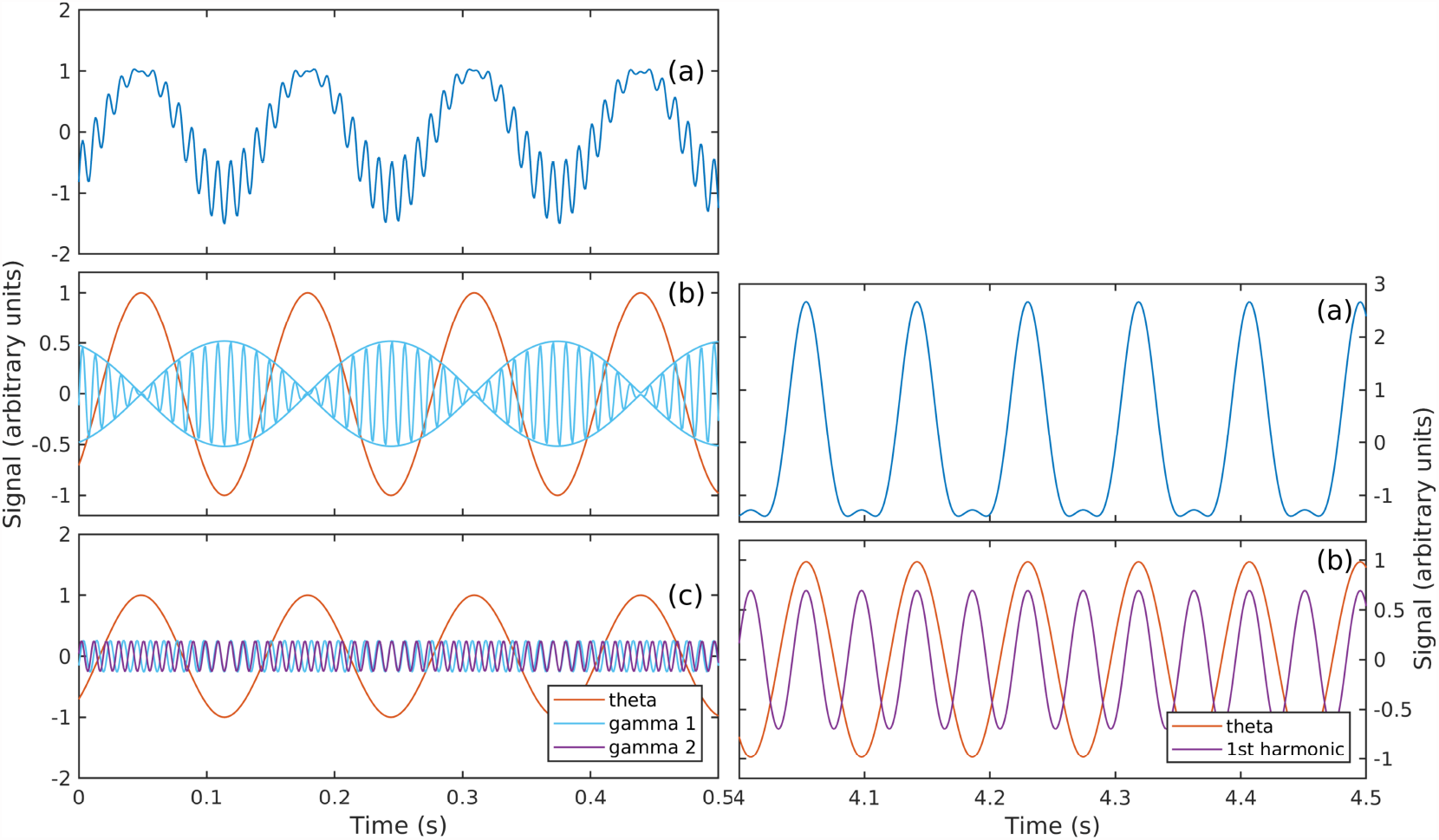
. Examples of numerical integration of equation for stationary states, under the assumption of resonance (equations 4 and 13), with *b_j_* satisfying equation 14. Left: Triad of the theta-gamma interaction type (e.g., figure 2) at stationarity, with 180° biphase;a) LFP signal;b) theta (red), gamma groups and gamma envelope (blue);c) theta (red) and gamma (blue-purple) rhythms. Right: Triad of the type theta-harmonic type at 0° biphase; a) LFP signal;b) theta (red) and its first harmonic (purple).

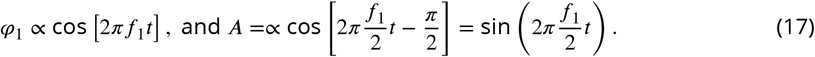

In other words, the gamma envelope is in quadrature with theta and its period is twice that of theta. This temporal structure is illustrated by integrating the three-wave model under the stationarity conditions defined by equations 4, 13, and 14, corresponding to 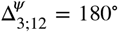. The gamma groups tend to be located in average in the troughs of theta, contributing overall negative skewness to the time series, consistent with the 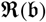 < 0 figure 4).

It is interesting to notice that the theta-harmonics type of interaction could possibly be captured by the Kuramoto description, as it agrees with the *m*Θ_1_ + *n*Θ_2_ = 0 paradigm;theta-gamma triad interaction, however, does not fit the paradigm and would likely be either aliased or missed altogether.

## Discussion

The present study was motivated by ambiguous results produced by efforts to detect the cross-scale integration that should be the hallmark of cognitive behavior (Hebb, 1958; Allen and Collins, 2013; Scheffer-Teixeira and Tort, 2016). The methodology used in previous phase-coupling studies is based on the fundamental assumption that the brain functions as a collection of independent oscillators that interact weakly. While the oscillators themselves are can be strongly nonlinear, they are assumed to be *essentially independent*, i.e., their interaction is weak enough that their phase-space orbit can be described as a “stiff limit cycle”: the interaction affects only their phase (“speed” along the periodic orbit).

This description does not seem applicable across the temporal scales of the hippocampal local-field potential. There is little observational support to the idea that LFP frequencies represent physical independent oscillators. Rather, recent observations (e.g., Lubenov and Siapas 2009; Patel et al. 2012) show that the hippocampal theta frequency band represents a propagating wave, i.e., a spontaneous neural activity organization that does not have a fixed physical oscillatory support. Perhaps more importantly, energy flow through the brain should also play an important part in scale integration, along with phase coupling. Phase models are, however, intrinsically equilibrium models: they can describe only phase rearrangements on top of a frozen scale distribution of energy.

Our observations of hippocampal LFP spectral show a strong correlation with both the rat speed and the hippocampal layer. They exhibit power law *f^−α^* behavior in at least two frequency bands (theta and gamma), with distinct slopes that also vary depending on the hippocampal layer. Remarkably, the spectral evolution as a function of rat speed shows synchronous increase in theta and gamma energy by orders of magnitude. The evolution of gamma in particular, showing the development of a front-like slope shifting toward higher frequencies, is strongly suggestive of a cross-scale energy flux from the theta frequency band. From the point of view of scale integration, the evolution of the scale-distribution of energy as a function of rat speed provides an important clue: nonlinear interactions are established as activity increases, and are intrinsically connected to the flow of energy.

This observation suggests the alternative description. The three-wave interaction model we propose as a paradigm for theta-gamma coupling is a universal conceptual model that has been studied intensely in other physics fields (Weiland and Wilhelmsson, 1977; Craick, 1985; Kartashova, 2010). This change of paradigm has important consequences.

As an immediate gain, the three-wave interaction model provides a fundamentally different theoretical platform for developing new analysis tools, and an alternative framework for the interpretation of observations. Because the theory is well established, the interpretation of the results is straightforward, and their statistics is well understood. An example is the use of phase and amplitude coupling estimators based on the bispectrum, e.g., 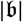. As shown in figure 4, bispectral estimates produce clear, unambiguous evidence of cross-spectral theta-gamma coupling, as well as its amplification in response to increasing rat speed. They also highlight the diversity of response of different hippocampus layers, some of which is seen for the first time and still to be understood.

The three wave interaction model suggests that the observations are consistent with quasi-stationary states, i.e., brain activity states in which the amplitude of different LFP rhythms is almost constant. While this seems to contradict our observations (and basic assumption) of significant evolution of brain activity, this contradiction is only superficial. The global state, measured here as a response to rat speed, evolves on a much longer time-scale (measured, say, in theta cycles) than LFP frequency bands, such that at every step of the way the frequency bands are allowed to relax in a “temporary equilibrium state”). This is consistent, for example, with the capacity of the rat to maintain a certain speed range for significant time intervals.

It is important to understand that the three-wave model is only a “toy” model, a conceptual model. Although, we believe, it provides valuable insight into the dynamics of the LFP components, it has strong limitations. For example, because the interactions described by the three-wave model cause only cyclic transfers of energy (the total energy is conserved if the system 5 is at resonance), therefore it does not describe how energy is transferred through the brain, and does not provide an explanation for the formation of the gamma front seen in the spectral evolution. To remove this limitation requires developing a model that describes the interaction within a full spectrum, not just a single triad. If the long-time, average behavior of a full-spectrum interaction is also taken into account, one needs to consider the scale and effect of energy sources and sinks. The total energy in a resonant triad is constant, but undergoes cyclic redistribution. If more triads of interacting modes are taken into account, and some modes participate in several triads, as is the case in a full spectrum, the energy transfers becomes more complicated, but the total energy is conserved. The system is in a stationary state. Imagine now that there is a triad involving a mode that leaks energy, i.e., transfers small amounts of energy *out of the system*. Even though in the leading order the exchange remains cyclical, a weak energy flow develops, directed toward the leaking *mode*. In this case a stationary state can be maintained only if energy is pumped into the system at the leakage rate. The theoretical framework for the study of such systems is known as “turbulence”, and the energy flow is usually referred to as the “scale cascade” (Richardson, 1922; Kolmogorov, 1941; Zakharov et al., 1992; Newell et al., 2001; Nazarenko, 2011). This theory might provide important information about the scale integration process. A discussion of a possible turbulence approach to the description of hippocampal activity is presented in a stochastic companion of this paper.

We conclude noting that, by approaching this problem from the physics perspective, it is possible to develop both kinetic and dynamic descriptions of these interactions with the intent of developing diagnostic tools as well as potential interventions. Our hope is that, ultimately, it may be possible to connect this approach with other formulations such as phase equations, or selforganized criticality, to describe the full scale integration brain in order to determine the physics of cognition.

## Materials and Methods

### Surgical Procedures

Surgery and all other animal care and procedures were conducted in accordance with the NIH Guide for the Care and Use of Laboratory Animals and approved by the Institutional Animal Care and Use Committee at the University of Florida. Rats were initially sedated in an induction chamber. Once anesthetized, the rat was transferred to a nose cone. The head was shaved with care taken to avoid the whiskers. The rat was then transferred to the stereotax, gently securing the ear bars and placing the front teeth over the incisor bar. The stereotaxic nose cone was secured, ensuring that the rat was appropriately inhaling the anesthesia. During surgical implantation, the rats were maintained under anesthesia with isoflurane administered at doses ranging from 0.5 to 2.5%. Next, ophthalmic ointment was applied and “tanning shades”, fabricated out of foil, were placed over but not touching the eyes to minimize direct light exposure. Multiple cycle of skin cleaning, using betadine followed by alcohol was applied prior to the first incision from approximately the forehead to just behind the ears. The remaining fascia was blunt dissected away and bone bleeding was mitigated through application of bone wax or cautery. Once the location of bregma was determined, the site of the craniotomy was located and a 3×3mm contour was drilled out, but not completed. This was followed by the placement of 7 anchor screws in the bone as well as a reference over the cerebellum and ground screw placed over the cortex. Once the screws were secured, a thin layer of dental acrylic (Grip Cement Industrial Grade, 675571 (powder) 675572 (solvent); Dentsply Caulk, Milford, DE) was applied taking care to not obscure the craniotomy location. Finally, the craniotomy location was completed, irrigating and managing bleeding as necessary once the bone fragment was removed. Next a portion of the dura was removed, taking care to avoid damaging the vessels and the surface of the neocortex. Small bleeding was managed with saline irrigation and gel foam (sterile absorbable gelatin sponges manufactured by Pharmacia & Upjohn Co, Kalamazoo, MI; a division of Pfizer, NY, NY). The probe implant coordinates targeted the dorsal hippocampus (AP: −3.2 mm, ML: 1.5 relative to bregma, DV: −3.7 to brain surface).

Once the probe was in place, the craniotomy was covered with silastic (Kwik-Sil, World Precision Instruments, Sarasota, FL) and then secured to the anchor screws with dental acrylic. Four copper mesh flaps were placed around the probe providing protection as well as acting as a potential Faraday cage. The wires from the reference and ground screws were soldered to the appropriate pins of the connector. Adjacent regions of the copper mesh flaps were soldered together to ensure their electrical continuity and the ground wire soldered to the copper mesh taking care to isolate the reference from contact with the ground. Once the probe was secured, the rat received 10cc of sterile saline as well as metacam (1.0 mg/kg) subcutaneously (the non-steroidal anti-inflammatory is also known as meloxicam; Boehringer Ingelheim Vetmedica, Inc., St. Joseph, MO). The rat was placed in a cage and monitored constantly until fully recovered. Over the next 7 days, the rat was monitored to ensure recovery and no behavioral anomalies. Metacam was administered the day following surgery as well. Antibiotics (Sulfamethoxazole/Trimethoprim Oral Suspension at 200mg/40 mg per 5 mls; Aurobindo Pharma USA, Inc., Dayton, NJ) were administered in the rat mash for an additional 5 days.

### Analysis and Statistics

Speed was calculated as the smoothed derivative of position. The local-field potential data was analyzed in Matlab^®^ (MathWorks, Natick, MA, USA) using custom written code as well as code imported from the HOSAtoolbox (Swami et al., 2001). Raw LFP records sampled at 24 kHz (Tucker-Davis system) were low-pass filtered down to 2 kHz and divided into sequences of 2048 time samples (approx. 1 second). The analysis of the LFP in the current study was based on standard techniques used for stationary signals (Priestley, 1981; Papoulis and Pillai, 2002) as previously described in Sheremet et al. (2016). Briefly, we assume that the LFP time series is a stochastic process, stationary in the relevant statistics, and decompose it using the discrete Fourier transform (DFT) The Fourier transform time sequences were reduced to the non-redundant frequency domain of 1 Hz ≤ *f* ≤ 1 kHz, with an frequency increment of 1 Hz.

Hippocampal layers were determined from the location of current sources and sinks derived on ripple and theta events, gamma power, and the polarity of the sharp-wave (Buzsaki, 1986; Buzsaki et al., 1986; Bragin et al., 1995; Ylinen et al., 1995; Lubenov and Siapas, 2009; Fernandez-Ruiz et al., 2017). Following an initial visual inspection the LFP was analyzed through bispectral analysis. Specifically, the nonlinearity of a time-series is expressed by the phase correlation across different frequencies. The lowest order (third order) phase coupling, described by the bispectrum and first used in ocean waves by Hasselmann et al. (1963), was used to analyze the hippocampal LFP as described in Sheremet et al. (2016). It is emphasized here as well as elsewhere (Van Milligen et al., 1995; Pradhan et al., 2012) that the bispectrum measures phase coupling, defined to occur when the sum of phases between two frequencies is equal to the value of a third frequency plus a constant. In order to associate our analysis with speed, the bispectrum for each channel are calculated across 1 second LFP fragment with mean speed of each segment is calculated respectively. Based on mean velocities of every segment, we classify each LFP segments into 4 speed ranges: 0.001 to 5 cm/s; 5 to 15 cm/s; 15 to 35 cm/s; and > 35 cm/s. For statistical comparisons, the 5 to 15 cm/s speed bin was compared to the > 35 cm/s bin.

## Acknowledgments

This work was supported by the McKnight Brain Research Foundation, and NIH grant MH109548. Special thanks to Sarah Lovett and Kim Robertson for technical support. AS thanks Prof. Victor Shrira for helpful discussions and suggestions.

## Appendix 1

### A Generic Nonlinear Model for Cross-frequency Coupling Formulation of the Problem

Here, we postulate that the brain activity on foliations can be represented as a multiple-scale, weakly-nonlinear wave field, whose dynamics is governed by nonlinear interaction, and present a simplified nonlinear model that exhibits nonlinear properties that agree with observational data. Forsimplicityand illustration purposes, instead of a discrete two-dimensional lattice, we confine the discussion to a one-dimensional spatial system whose state is defined by the function *ϕ*(*x,t*), e.g., electric potential, of spatial coordinate vector *x* and time *t*. Denote the deviation of *ϕ* from a reference state *ϕ*_0_ by *φ* = *ϕ* – *ϕ*_0_. If deviation remains small at all times, i.e., *φ* = *O*(*εϕ*), with *ε* ≪1, under quite general conditions *φ* satisfies a nonlinear equation of the form

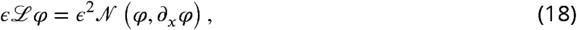

where 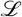 is a linear differential operator, and 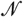 function nonlinear in its argument. The function 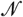 will be assume here to be a polynomial of second degree (hence the scaling factor *ε*^2^) to represent a “minimum” nonlinearity.

### Linear system

In the limit *ε* → 0 equation becomes linear,

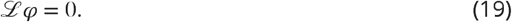

and a standard approach to solve it is to use the Fourier transform

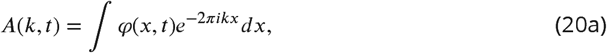

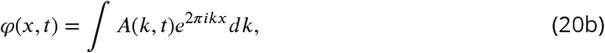

where *k* = 1/*λ* is the wave number with *λ* wavelength, and functions *e^2πikx^* are orthogonal in the sense that

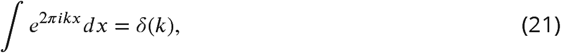

with δ the Dirac delta function. In equation 20b, *φ* is real if *A*(*-k,t*) = *A*\*(*k,t*). Setting

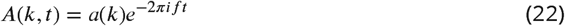

where *f* = 1/*T* is the frequency (*T* is the wave period), and substituting the decomposition 20b into equation 19 yields the dispersion relation Whitham (1974)

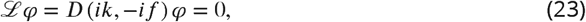

an algebraic equation/expression relating *f* and *k*. The dispersion relation defines the connection between temporal and spatial scales (Buzsaki and Draguhn, 2004), and the linear properties of the system: phase and group speeds, and conservation of energy. If the neural lattice is finite the Fourier space is discrete. The discretization of the Fourier space may be achieved formally by sampling *A*(*k, t*) at discrete values {*k_j_*}_*j=1,…,n*_ i.e., writing

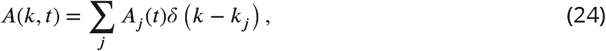

where *δ (k)* is Dirac’s delta function, which yields the general solution in the form

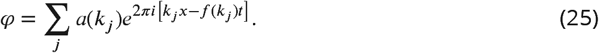

The linear system illustrates the duality of wave-number and frequency representations. The structure of the solution of the linear system is determined by two parameters, *k* and *f*, but due to the dispersion relation constraint 23, one of the two parameters is free. The choice of the free parameter is arbitrary, and provides some freedom in formulating the dynamics. Setting the wave number as independent parameter implies that the spatial structure is well resolved and the equation describes the evolution of the amplitudes in time, c.f. equation 27. Setting the frequency as the independent parameter implies that the solution is stationary; the resulting equation describes the evolution in space. The latter approach is better suited for the analysis of observations obtained at a single point in space; the former approach is the natural physical (future is unknown), and also better suited for boundary value problem, where the spatial structure is dictated by the boundary conditions. Below, we assume the spatial structure known: *k* is the free parameter and, for simplicity, the dispersion relation 23 has a single root (mode) *ω = *ω*(k)*.

### Dynamical equation with quadratic nonlinearity

Because nonlinearities will force the solution to deviate from the harmonic form 22, in the nonlinear case (*ε* ≠ 0) solutions of equation 18 should be sought in the general form

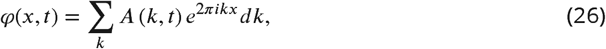

with no additional assumptions about the form of the time evolution. Substituting 26 into equation 18 yields the time evolution equation for the complex amplitudes, also called the dynamical equation

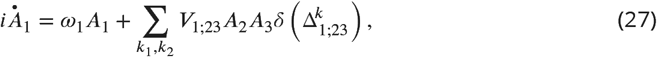

where 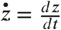, with *z* some variable, and with the shorthand notations *A_j_* = *A* (*k_j_, t*),) *ω_j_* = *ω*(*k_j_*), and 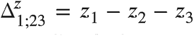, e.g., *k* in equation 27. In equation 27, the time evolution of the modal amplitude has two components: the linear component with radian frequency *ω*, and the contribution of the nonlinear interactions. The integral on the right-hand side represents nonlinear (triad) contributions to the evolution of *A*_1_. The interaction coefficient *V*_1;23_ is a function of the wave numbers *k_j_*, *j* = 1,2,3. The delta function *δ* acts as a selection criterion: only triads satisfying the equation *k*_1_ - *k*_2_ - *k*_3_=0 interact (i.e., are counted in the integral).

### The three-wave equation

The simplest non-trivial form of the dynamical equation 3 is obtained for a single triad of modes *k_j_*, *j* = 1,2,3, that satisfy the condition *k*_1_ + *k*_2_ − *k*_3_ = 0 (otherwise the sum in the right-hand side of equation 3 cancels). The simplified form is called the three-wave equation The discussion below is based on the texts by(Weiland and Wilhelmsson, 1977; Craick, 1985; Rabinovich and Trubetskov, 1989). With the substitutions 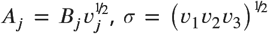, with *v*_1_ = *2V*_1;-23_, *v*_2_ = *2V*_2;-1,3_, and *v*_3_ = *2V*_3;12_, the equations for a triad acquire a standard convenient form

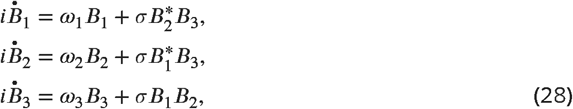

Multiplying equations 28 by 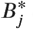 and subtracting their complex conjugate yields the equations

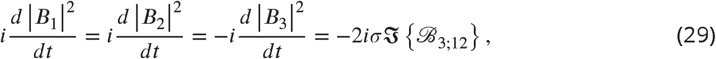

where 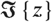 is the imaginary part of the complex number *z*. If model energy is defined as the amplitude square |*B_j_*|^2^, the system 29 describes the evolution of the modal energy. Note that in equation 29, the rate of change of modal energy depends on the triple product

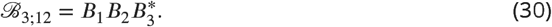

Switching to amplitude-phase representation *B_j_* = *b_j_e^iΘj^*,with *b_j_*, 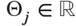 and *b_j_* >0, one can rewrite equations 28 as a system of equations for amplitudes

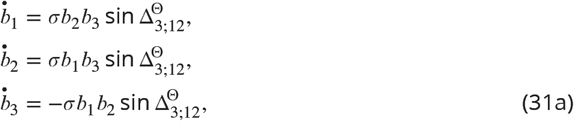

and one for the phases

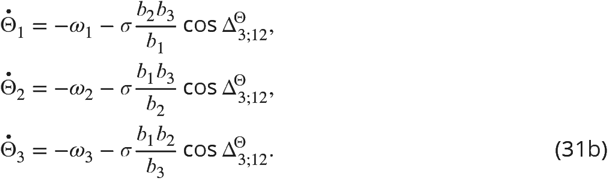

### Qualitative behavior of the 3-wave equations

The system 28 may be reduced to 4 equations and has analytical solution given in terms of Jacobi elliptic functions. Combining the last three equations yields a system of four equations with four unknowns: the amplitudes *b_j_*, *j* = 1,2,3, and the phase 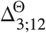

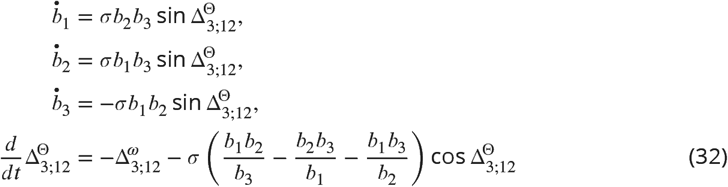

The system 32 has the Manley-Rowe integrals of motion

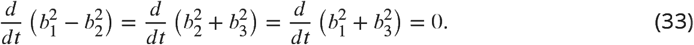

(note that only two of these relations are independent). Also, through direct differentiation one can also verify that the quantity

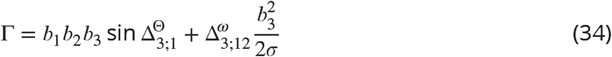

is conserved, providing the fourth conservation law 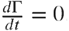.

The procedure for integrating the conservation laws: the Manley-Rowe relations can be integrated directly to yield

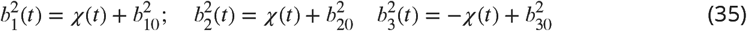

where *b_j0_* is the initial value of amplitude *b_j_*, and

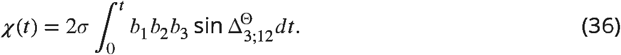

Combining the equations 35 and 34 yields

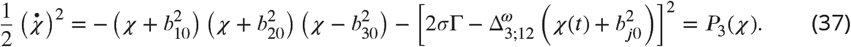

where *P*_3_ is a cubic polynomial in the unknown function *χ*.

The solutions of equation 37 is given in therms of Jacobi elliptic functions (Craick, 1985; Kartashova, 2010). Without going into the details of the analytical solutions, a qualitative understanding of the behavior of the solutions can be derived from the examination of the Manley-Rowe integrals (e.g. Rabinovich and Trubetskov 1989). If most of the energy of the triad is stored initially in the lowest wave number (or frequency), the energy transferred to higher wave numbers is small. Suppose that initially 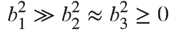 and that 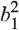 decreases over *dt* by the amount 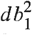. Equations 33 show that 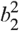 and 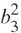 have to decrease/increase by the same amount 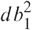; However, because 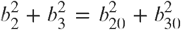 is small, 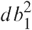 cannot exceed the minimum of 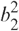 and 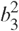 and has to change sign. The consequence is that the higher wave-number modes of a triad cannot grow at the expense of the lowest frequency mode. This behavior is known as decay instability. If, however, the highest wave-number mode is the most energetic, 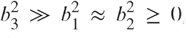, equations 33 show that both modes 1 and 2 grow simultaneously and the entire energy stored in mode 3 is distributed to 1 and 2. Because the evolution of the triad is determined significantly by the mode that initially stores most of the energy of the triad, the with the highest wave number (frequency) is called “active” and the rest of the modes are called “passive” (Kartashova, 2010).

## Footnotes

1 In Kuramoto’s words, “(the) limit cycle (of each oscillator) is supposed to possess more or less ‘stiffness’ in orbital shap against perturbations, so that when only weakly perturbed, the state point of a given oscillator hardly deviates from its natura closed orbit but remains almost on it. But the deviation in phase produced along the natural orbit could accumulate, thu needing a proper description. Since this argument applies to any oscillator, we are led to the following picture: the state of each oscillator can be approximately specified by its phase value, and its rate of change is determined by the phase values of all the other oscillators interacting with it.” (Kuramoto, 1984).

